# Albumin inhibits the activation of hepatic stellate cells by suppressing TGF-β/Smad3 signaling via IL-1β

**DOI:** 10.1101/753152

**Authors:** Ji Hoon Park, Janghyun Kim, So-Young Choi, Kiweon Cha, Heekyung Park, Jung-Eun Lee, Boram Lee, Ji Wook Moon, Sun-Hwa Park, Jae Min Lee, Hong Sik Lee, Junseo Oh

**Author notes:** Corresponding author: J Oh; Department of Biomedical Sciences, Korea University Graduate School, Seoul 02841, Korea. These authors contributed equally to this work (Ji Hoon Park, Janghyun Kim).

## Abstract

Activated hepatic stellate cells (HSCs) play a key role in liver fibrosis and inactivating HSCs has been considered a promising therapeutic approach. We previously showed that albumin and its derivative, retinol binding protein (RBP)-albumin domain III fusion protein (named R-III), inhibit HSC activation. Here, we investigate the mode of action of albumin and R-III. NF-κB in activated HSCs was evenly distributed in the cytoplasm, but albumin expression and R-III treatment (albumin/R-III) induced NF-κB nuclear translocation via retinoic acid (RA) sequestration, resulting in increased expression of interleukin-1β (IL-1β). In an IL-1β dependent manner, albumin/R-III inhibited Smad3 nuclear translocation via TAK1-, JNK-mediated Smad3 linker phosphorylation and decreased expression of Smad3 target genes, such as α-smooth muscle actin and collagen type I. Mutation of the Smad3 linker phosphorylation sites abolished R-III effects on Smad3. In conclusion, our data suggest that the anti-fibrotic effects of albumin/R-III are due to RA sequestration which downregulates RAR-mediated signaling and also TGF-β/Smad3 signaling. This mechanistic elucidation of albumin function in HSCs provides clues to understanding the frequent albumin mutations found in hepatocellular carcinoma.

## Introduction

Stellate cells, commonly known as hepatic stellate cells (HSCs), are first mentioned by Von Kupffer in 1876 [1]. HSCs are pericytes found in the perisinusoidal space of the liver and constitute approximately 5-8% of total liver cells [2]. They are in a quiescent state with non-proliferative characteristic in the normal liver and represent the major storage site for retinoids. About 80 % of the total body vitamin A (retinol) is stored as retinyl esters in lipid droplets in the cytoplasm. Upon response to fibrogenic stimuli, quiescent stellate cells undergo functional and phenotypical changes, referred to as “activation”, and transform into myofibroblast-like cells [3]. This process is characterized by loss of cytoplasmic vitamin A-containing lipid droplets, high cellular proliferation, positive staining for α-smooth muscle actin (α-SMA), and greatly increased synthesis of extracellular matrix (ECM) proteins. It is widely accepted that the activation of HSCs plays a critical role in liver fibrosis, which is characterized by excessive deposition of extracellular matrix components [4]. Similar cells in pancreas were isolated in the late 1990s [5], and these pancreatic stellate cells (PSCs) also play an important role in pancreatic fibrogenesis in a manner analogous to HSCs [6]. Thus, stellate cells are considered as an attractive target for anti-fibrotic therapies [7]. Despite extensive investigations, there is, however, no effective therapy for liver fibrosis and end stage cirrhosis, except for the removal of the causative agent and organ transplantation.

Retinoids (vitamin A and its metabolites) regulate multiple physiological activities, such as vision, cell proliferation and differentiation [8]. Vitamin A, acquired from the diet, is transported to the liver and taken up by hepatocytes as a chylomicron remnant. It has been suggested that retinol binding protein (RBP) plays a role in the transfer of retinol from hepatocytes to HSCs via a RBP receptor STRA6 [9]. Upon HSC activation, some retinoid contents are oxidized to retinaldehyde, which is irreversibly further metabolized to retinoic acid (RA) by retinaldehyde dehydrogenases (RALDH). This is supported by the fact that RA level is increased in activated stellate cells compared with pre-activated stellate cells, whereas the contents of retinyl ester and retinol decrease [10, 11]. In order to examine the role of RA in stellate cell activation, studies have undergone by assessing the effects of exogenous RA on HSCs or liver fibrosis, but the results are controversial; several studies showed that RA inactivated HSCs and alleviated hepatic fibrosis [12, 13], while other reports showed the opposite [14, 15]. Studies showed that retinoids exert their biological effects primarily through binding to nuclear receptors, retinoic acid receptors (RAR) and retinoid X receptor (RXR) [16].

Albumin is an abundant multifunctional plasma protein synthesized primarily by liver cells [17]. It, comprised of three homologous domains (I-III), binds a wide variety of hydrophobic ligands including fatty acids, retinoids [18, 19]. In a previous study, we showed that albumin was endogenously expressed in quiescent but not activated stellate cells and that its forced expression in activated stellate cells induced phenotypic conversion of myofibroblasts into fat-storing cell phenotype [20]. Based on this finding, we have developed the recombinant fusion protein R-III, in which albumin (domain III) is fused to the C-terminus of RBP [21]. RBP was adopted for stellate cell-targeting delivery because RBP and its membrane receptor STRA6 coordinate cellular uptake of retinol into HSCs [9, 22]. Our follow-up study showed that R-III inhibits HSC activation *in vitro* and reduces liver fibrosis *in vivo* [23, 24]. In this study, we examined the mode of action of albumin/R-III and found that RA sequestration by albumin/R-III downregulates TGF-β/Samd3 signaling in HSCs.

## Results

### Retinoic acid is involved in the activation of hepatic stellate cells (HSCs)

Retinoic acid (RA) levels were reported to increase in activated HSCs [10], and we previously showed that the inhibition of HSC activation by albumin expression and R-III treatment (albumin/R-III) was accompanied by a pronounced reduction of RA levels and its signaling [23]. In order to examine the role of RA in HSC activation, HSCs after passage 1 (HSCs-P1; activated HSCs) were treated with citral, a RALDH inhibitor that blocks RA biosynthesis, or AGN193109, a RAR antagonist, and analyzed by real-time PCR. Citral significantly downregulated the expression of α-SMA and collagen type I, markers for activated stellate cells, whereas AGN193109 affected only α-SMA expression (Fig. 1A). This suggests that RA is involved in HSC activation through as yet unidentified pathway(s) rather than via its nuclear receptor RAR.

**FIG. 1.**
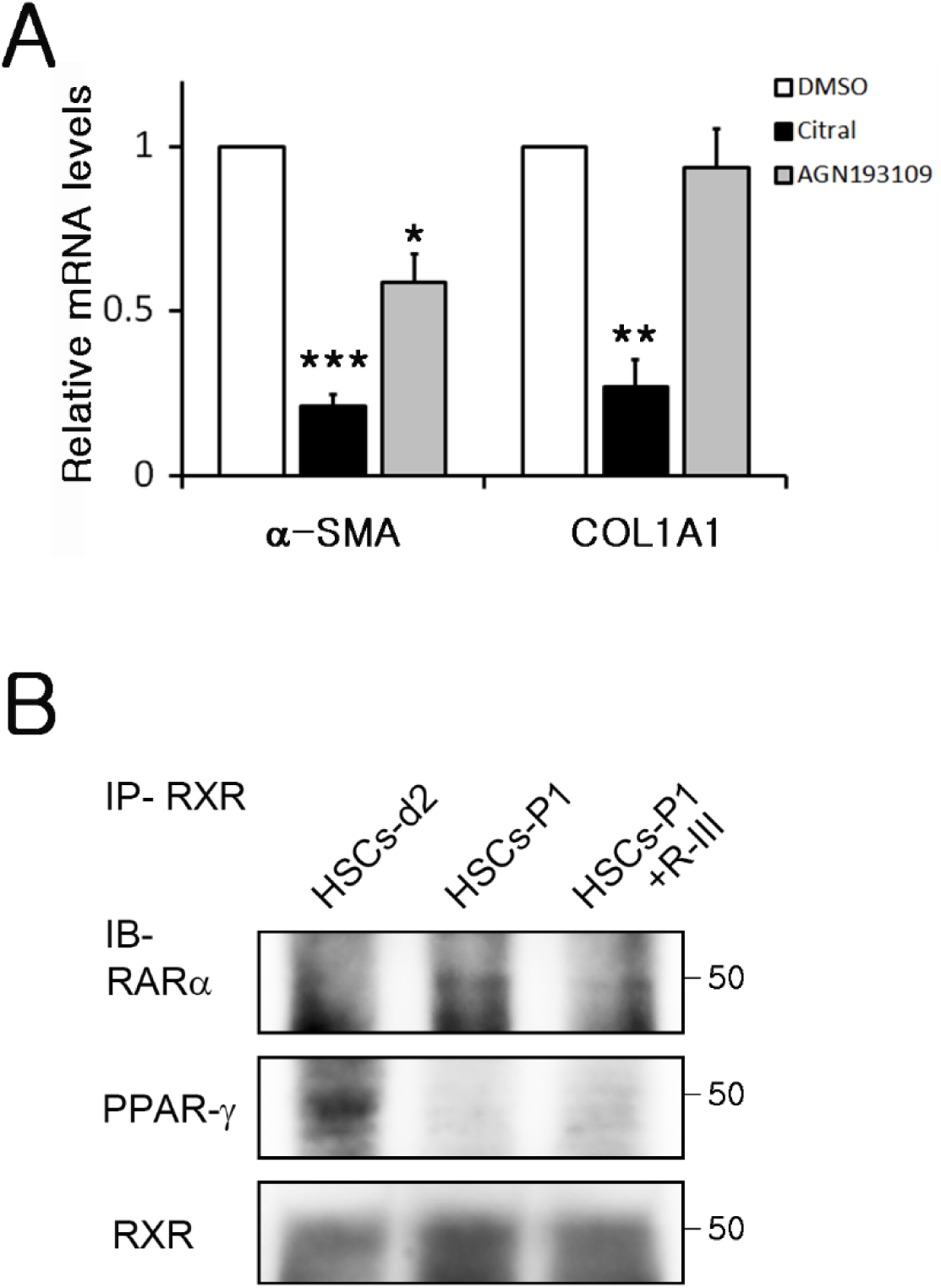
Retinoic acid is involved in HSC activation. (A) HSCs after passage 1 (HSCs-P1) were treated with citral (100 μM) or AGN193109(1 μM) for 20 h and analyzed by real-time PCR for the expression levels of α-SMA and collagen type I. The PCR data are expressed as the ratio of untreated control and represent the means (SD) for three independent experiments. *P*-value, paired t-test (compared to untreated HSCs-P1): *P < 0.05, **P < 0.01, ***P < 0.001 (B) HSCs at day 2 after plating (HSCs-d2), HSCs-P1 and R-III-treated HSCs-P1 were immunoprecipitated with anti-RXR antibody and subject to western blotting. The Western blots are representative of two independent experiments from separate cell preparations.

Next, we examined how the alterations in RA levels influence nuclear receptor signaling. Cell lysates were prepared from early-activated HSCs (HSCs at day 2 after plating; HSCs-d2), HSCs-P1 and R-III-treated HSCs-P1, respectively, and immunoprecipitated using antibody against RXR, an obligatory heterodimerization partner. In parallel with RA levels reported, the heterodimer formation of RXR with RAR-α was found to increase in activated HSCs compared with early-activated HSCs and decrease with R-III treatment (Fig. 1B). On the other hand, the heterodimer formation of RXR with PPAR-γ, a factor whose activity is decreased in activated HSCs and whose agonist suppresses several markers of HSC activation [25], diminished in activated HSCs and remained undetectable even with R-III treatment. This suggests that R-III partially reverses HSC activation.

### RA sequestration is accompanied by nuclear translocation of NF-κB in HSCs

The NF-κB family of transcription factors are central players in inflammatory response, and they are normally sequestered in the cytosol by inhibitor κB (IκB) and upon activation translocate to the nucleus [26]. As it was reported that RA exerts an antiinflammatory effect by inhibiting nuclear translocation of NF-κB in microglial and neuronal cells [27, 28], we assessed the subcellular localization of NF-κB P65 in HSCs by immunofluorescence. P65 were found distributed evenly in the cytoplasm in activated HSCs, and, intriguingly, albumin/R-III led to a strong nuclear translocation of p65 (Fig. 2). Such p65 nuclear translocation was not, however, detected in HSCs-P1 expressing mutant albumin (R410A, Y411A, K525A), in which three high affinity-fatty acid binding sites were substituted for alanine and whose expression failed to sequester RA [23]. This suggests that RA in activated HSCs may prevent nuclear import of NF-κB p65. Indeed, RNA-seq analysis revealed that R-III treatment increased the expression of inflammation-related genes, such as IL-1α, IL-1β, IL-6 and TNF-α, in HSCs-P1 (Table 1). Real-time PCR also confirmed that IL-1β expression was dramatically elevated by albumin/R-III by >100-fold (data not shown).

**Table 1.** Expression profile of inflammation-related genes

| Gene | HSCs-d5/HSCs-d2 | R-III-treated HSCs-P1/HSCs-P1 |
| --- | --- | --- |
| Interleukin 1α | -7.13 | +18.67 |
| Interleukin 1β | -7.52 | +135.18 |
| Interleukin 6 | -4.46 | +4.24 |
| TNF-α | -3.63 | +2.57 |

Total RNA was extracted from HSCs at day 2, 5 after plating (HSCs-d2, HSCs-d5) and from HSCs after passage 1 (HSCs-P1) and R-III-treated HSCs-P1, and analyzed by RNA sequencing.

**FIG. 2.**
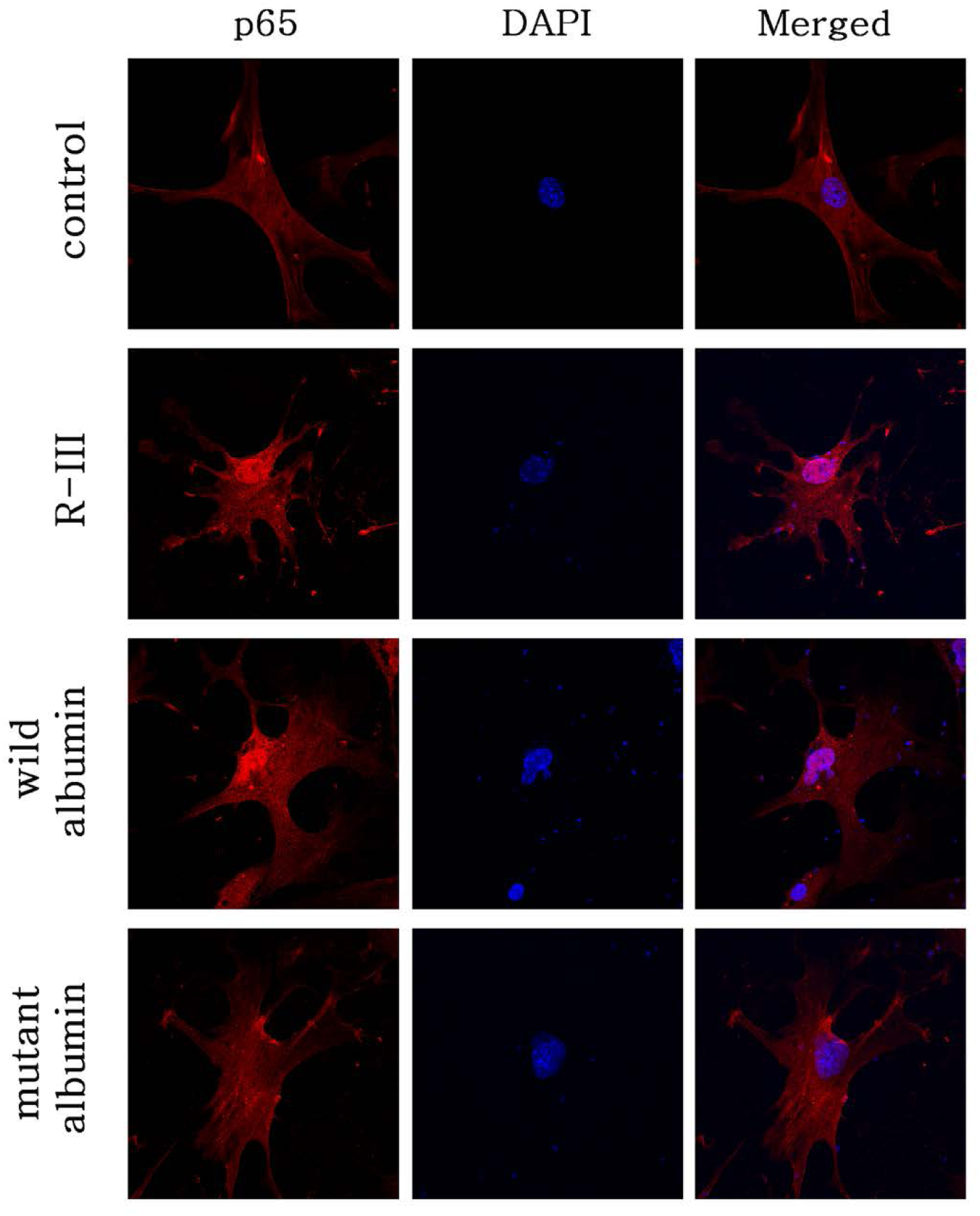
R-III treatment and albumin expression promote nuclear translocation of NF-κB. HSCs-P1 were treated with HPLC-purified R-III (0.3μM) or transfected with plasmid encoding wild-type or mutant albumin (R410A/Y411A/K525A), and analyzed by immunofluorescence using anti-p65 antibody. The cell nuclei were stained with DAPI (blue).

### IL-1β mediates R-III effects on HSC activation

Previous reports showed that IL-1β neutralizes the activity of TGF-β, a major driver of HSC activation, by different mechanisms, such as TGF-β receptor type II (TGF-βRII) downregulation, Smad7 upregulation, and Smad3 inhibition [29-31]. We first examined whether IL-1β influences HSC activation. Surprisingly, IL-1β expression in activated HSCs downregulated the mRNA expression of α-SMA and collagen type I, as was seen with albumin expression and R-III treatment [20] (Fig. 3A). We then tested whether R-III effects on α-SMA and collagen type I is mediated by IL-1β and found that co-treatment with IL-1 receptor antagonist (IL-1RA) abolished R-III effects (Fig. 3B). The time course of changes in mRNA levels of IL-1β, α-SMA and collagen I was examined with R-III treatment. R-III induced a steady increase in IL-1β mRNA with a peak in expression at ∼12 h (>100 fold) and simultaneously downregulated both α-SMA and collagen I (Fig. 3C). Lastly, we investigated R-III effect on the mRNA levels of TGF-βRII, smad7 and TGF-β1, which are listed as TGF-β/Smad3 target genes [32], and found modest but significant decreases in Smad7 and TGF-β (Fig. 3D). Thus, these findings suggest that RA sequestration by albumin/R-III triggers nuclear import of NF-κB and that the resulting IL-1β may mediate R-III effects on HSC activation.

**FIG. 3.**
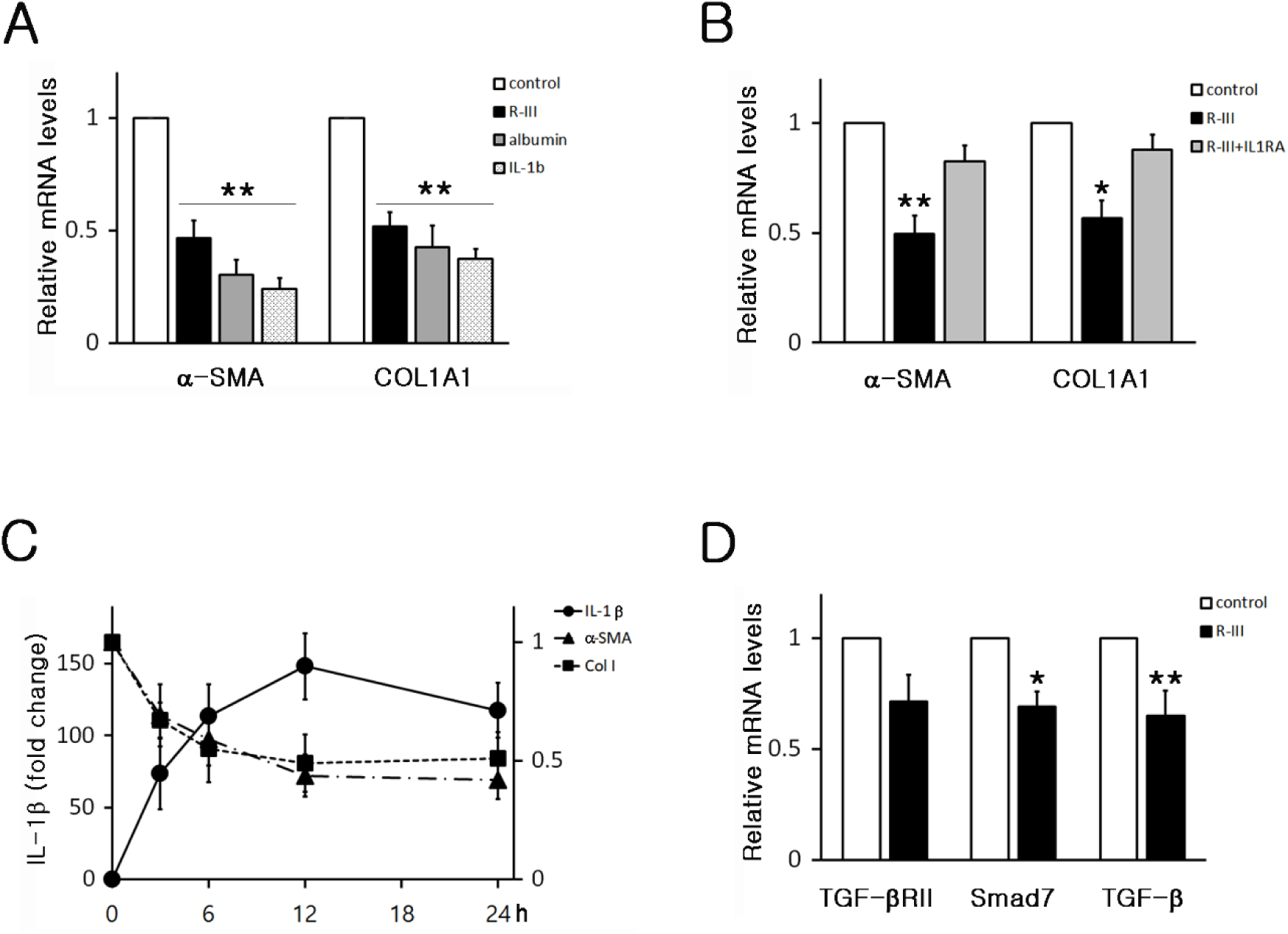
The inhibitory effects of R-III on HSC activation is mediated by IL-1β. (A) HSCs-P1 were treated with R-III (0.3 μM) for 20 h or transfected with plasmid encoding albumin, R-III and IL-1β and analyzed by real-time PCR for the expression levels of α-SMA and collagen type I. (B) HSCs-P1 were treated with R-III in the presence or absence of IL-1RA (1 μg/ml) and analyzed by real-time PCR. (C) HSCs-P1 were treated with R-III, harvested at the indicated time points and analyzed by real-time PCR for the expression levels of IL-1β, α-SMA and collagen type I. Fold changes of IL-1β mRNA (left y-axis) and of α-SMA and collagen I mRNA (right y-axis). (D) HSCs-P1 were treated with R-III and analyzed for the expression levels of TGF-β receptor type II, Smad7 and TGFβ1. *P*-value, paired t-test (n = 3) (compared to untreated HSCs-P1), *P < 0.05, **P < 0.01

### Albumin/R-III inhibits nuclear translocation of smad3

As IL-1β was reported to affect subcellular distribution of Smad proteins [31], we tested whether increased production of IL-1β by albumin/R-III inhibits nuclear translocation of Smads. Immunofluorescence showed that both Smad2 and Smad3 were found accumulated in the nucleus in activated HSCs. Albumin/R-III was, however, found to block the nuclear import of smad3, but not Smad2, in a IL-1β dependent manner (Fig. 4A and 4B). As importin7 and 8 are reportedly responsible for transporting phosphorylated Smad2 and Smad3 into the nucleus [33], we measured their expression levels but found no significant changes with R-III treatment (Fig. 4C).

**FIG. 4.**
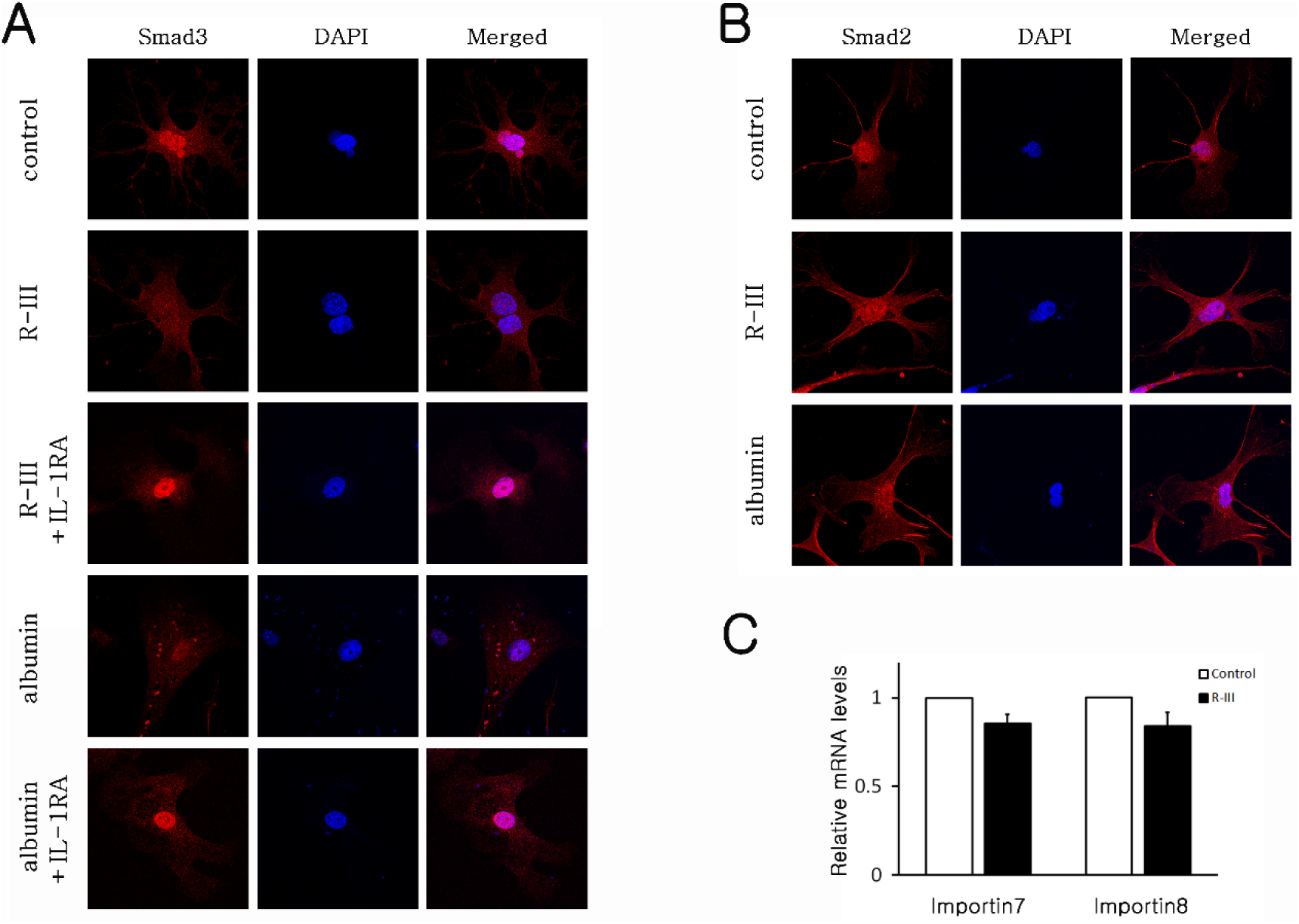
R-III treatment and albumin expression inhibits nuclear translocation of Smad3 in HSCs. (A) HSCs-P1 were treated with R-III (0.3 μM) or transfected with albumin expression vector, in the presence or absence of IL-1RA, and analyzed by immunofluorescence using antibodies against Smad3. (B) Immunofluorescence images using anti-Smad2 antibody. (C) HSCs-P1 were treated with R-III and analyzed by real-time PCR for the expression levels of importin7 and 8.

### IL-1β induces Smad3 linker phosphorylation

TGF-β-activated kinase 1 (TAK1) and its downstream targets, p38 and JNK, have been identified that phosphorylate Smad3 in the linker region and inhibit its nuclear translocation [34, 35]. Other studies also demonstrated that IL-1β inhibits TGF-β signaling through Smad linker phosphorylation in a TAK1-dependent manner [31, 36]. We examined whether albumin/R-III modulates the kinase activities and Smad3 linker phosphorylation via IL-1β in HSCs by western blot analysis. As it was reported that phosphorylation of TAK1 at Ser412 is required for activation of TAK1-mediated IL-1R signaling [37], TAK1 S412 phosphorylation was increased by albumin/R-III (Fig. 5A, Supplementary Fig. S1), but no significant change in phosphorylation at TAK1 autophosphorylation sites T184, T187 was observed. Importantly, albumin/R-III activated JNK, but not p38, and enhanced phosphorylation of Smad3 in the linker region (S208). Smad3 phosphorylation at the C terminus S423, S425 was not affected. These R-III effects on Smad3, JNK and α-SMA were found to be dependent on IL-1β signaling (Fig. 5B). In order to identify the kinase(s) responsible for Smad3 linker phosphorylation, we co-treated HSCs-P1 with R-III and the inhibitor for p38 (SB203580), JNK (SP600125) or TAK1 ((5Z)-7-Oxozeaenol) and found that Smad3 linker phosphorylation was reduced by (5Z)-7-Oxozeaenol and SP600125 (Fig. 5C). Real-time PCR also showed that the activity of TAK1 and JNK was required for R-III effects on α-SMA and collagen type I, TGF-β/Smad3 target genes (Fig. 5D). Lastly, in order to confirm the functional significance of Smad3 linker phosphorylation, we transfected HSCs-P1 with expression vector for wild-type or mutant Smad3 (S204A, S208A, S213A), in which three linker phosphorylation sites were substituted for alanine. R-III effects on the subcellular localization of Smad3 and on the expression of α-SMA and collagen I were not detected in mutant Smad3-expressing HSCs (Fig. 6A and 6B). These findings suggest that IL-1β signaling induced by albumin/R-III inhibits nuclear import of Smad3 by linker phosphorylation.

**FIG. 5.**
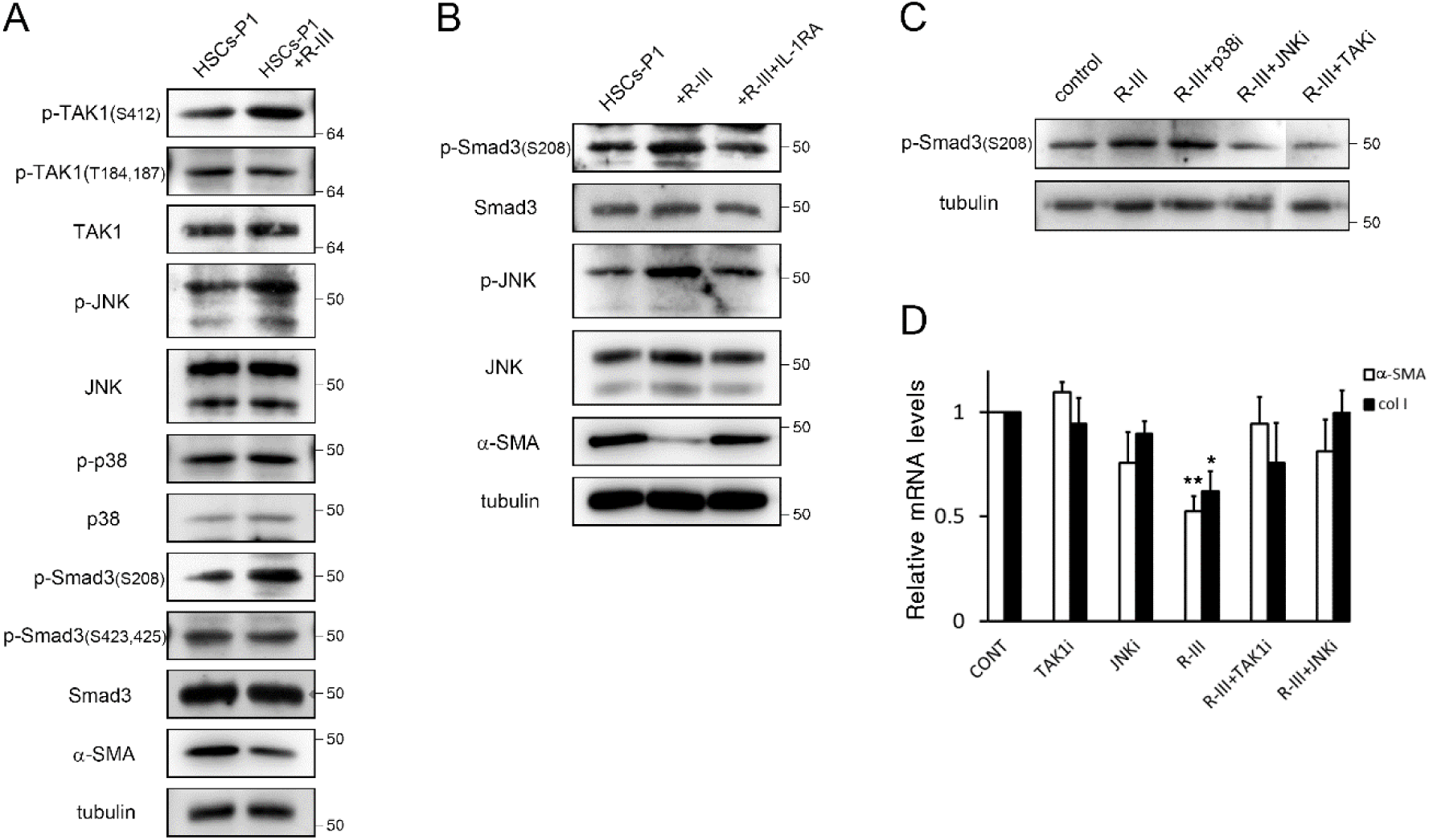
IL-1β induces linker phosphorylation of Smad3 in HSCs. (A) HSCs-P1 were treated with R-III (0.3 μM) for 20 h and cell lysates were analyzed by western blotting. The Western blots are representative of three independent experiments from separate cell preparations. α-tubulin was used as a loading control. (B) HSCs-P1 were treated with R-III in the presence of IL-1RA (1 μg/ml) and analyzed by western blotting. (C, D) HSCs-P1 were treated with R-III in the presence of the inhibitor for TAK1 ((5Z)-7-Oxozeaenol, 0.25 μM), p38 (SB203580, 5 μM) and JNK (SP600125, 10 μM) and analyzed by western blotting (C) and real-time PCR (D). *P*-value, paired t-test (n = 3) (compared to untreated HSCs-P1), *P < 0.05, **P < 0.01

**FIG. 6.**
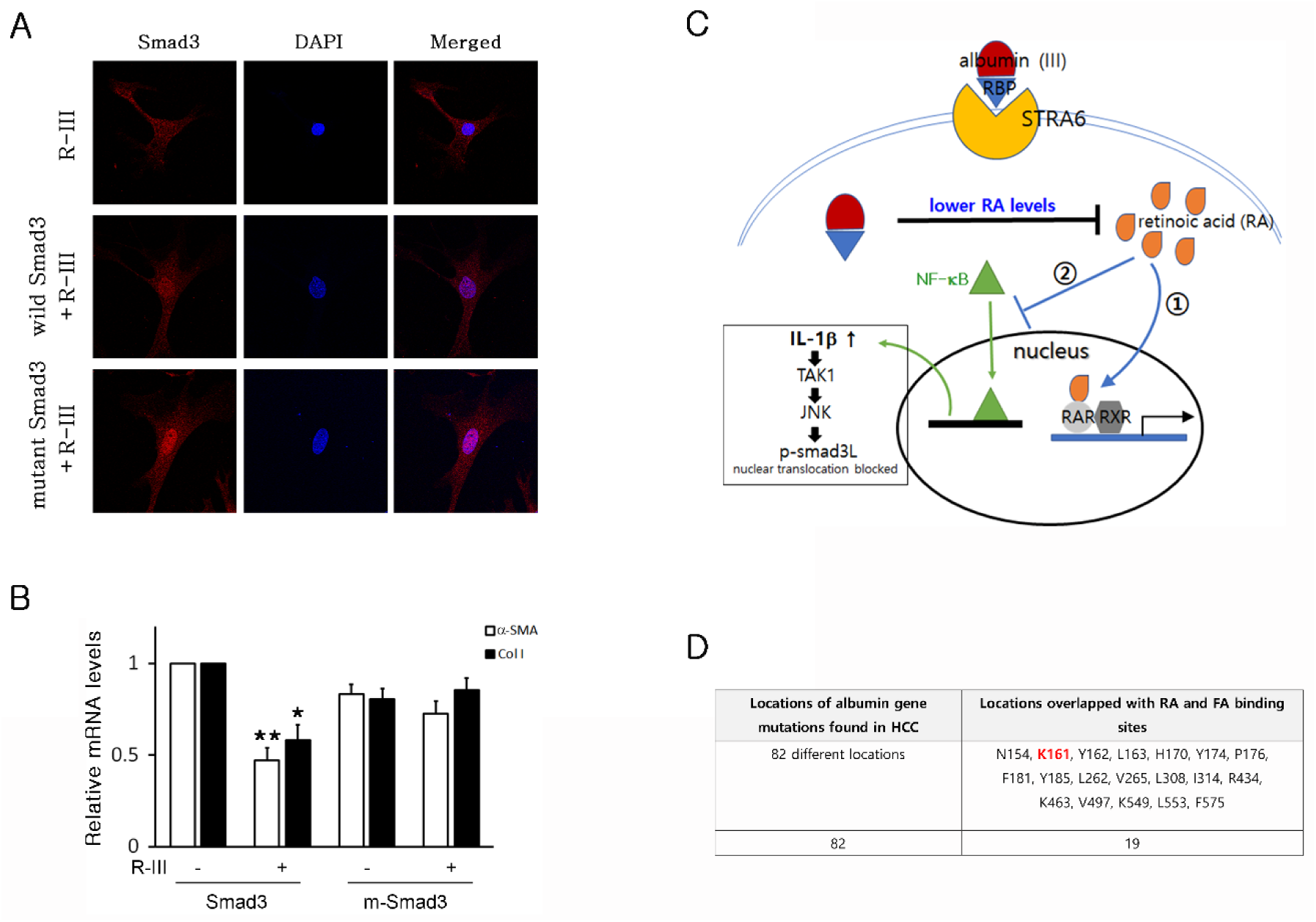
Smad3 linker phosphorylation is important for the anti-fibrotic effects of R-III. (A, B) HSCs-P1 were transfected with plasmid encoding wild-type or mutant Smad3(S204A, S208A, S213A), treated with R-III (0.3 μM) for 20 h and analyzed by immunofluorescence using anti-Smad3 antibody (A) or real-time PCR (B). *P*-value, paired t-test (n = 3) (compared to untreated HSCs-P1), *P < 0.05, **P < 0.01 (C) Schematic drawing of the mechanism of action of R-III in HSCs. R-III enters HSCs via a RBP receptor STRA6 and sequesters RA, which not only ➀ blocks RAR signaling but also ➁ inhibits Smad3 nuclear translocation via NF-κB/IL-1β action. (D) The locations of albumin mutations found in HCC overlap binding sites for RA and fatty acid (FA).

### Albumin mutation may be associated with hepatocellular carcinoma

As our finding suggests that RA sequestration by albumin/R-III downregulates not only RAR-mediated signaling but TGF-β/Smad3 signaling, leading to the inhibition of HSC activation (Fig. 6C), it is plausible to reason that mutations in the albumin gene, particularly at the sites that are involved in RA binding, may impair its anti-fibrotic activity in HSCs and predispose a person to fibrosis, cirrhosis and liver cancer, as cirrhosis is the major risk factor of hepatocellular carcinoma (HCC). Intriguingly, albumin gene was recently reported to be frequently mutated in liver cancer [38]. Among 82 locations of albumin gene mutations found in HCC, publicly available at www.cbioportal.org [39], 19 overlapped binding sites for RA and fatty acid [18, 19] and, more importantly, the most frequently mutated amino acid (K161) was found to be a RA binding site (Fig. 6D). RA binds also to the albumin fatty acid binding sites [40]. This high degree of overlap reinforces the anti-fibrotic role of albumin in HSCs.

## Discussion

Hepatic stellate cells are considered an attractive target for anti-fibrotic therapies. Several previous reports aimed to inactivate HSCs and reduce liver fibrosis, but no effective therapy for treating liver fibrosis is currently available [41]. Also the molecular mechanism for the activation of HSCs remains elusive despite intensive research efforts over the last 30 years.

Quiescent stellate cells store retinoids as retinyl esters in cytoplasmic lipid droplets. Upon stellate cell activation, cytoplasmic lipid droplets collapse and a portion of the retinoid contents is likely released and metabolized to retinaldehyde and to retinoic acid (RA). The role of RA in HSC activation has been proposed, but previous reports about the effects of exogenous retinoids on HSCs and liver fibrosis were controversial [42]. On the other hand, we previously showed that endogenous RA may be involved in HSC activation [23], and demonstrated in this study that RA inhibits NF-κB nuclear transport in HSCs (Fig. 2). Consistent with the previous finding that albumin expression and R-III treatment (albumin/R-III) sequestered cellular RA in HSCs [23, 24], albumin/R-III allowed nuclear translocation of NF-κB (Fig. 2) and led to a dramatic increase in IL-1β expression. IL-1β signaling in turn activated TAK1 and its downstream molecule JNK (Fig. 5A), phosphorylated Smad3 in the linker region, inhibiting nuclear transport of Smad3 (Fig. 4A). As a result, albumin/R-III in HSCs downregulated the expression of Smad3-target genes, such α-SMA, collagen type I, TGF-βRII, Smad7, and TGF-β. This mode of action for albumin/R-III was further supported by the finding that mutation of the Smad3 linker phosphorylation sites (S204A, S208A, S213A) abolished R-III effects on Smad3 (Fig. 6A). Thus, the anti-fibrotic effects of albumin and R-III is probably due to reduction in cellular RA levels. This explains the reason why mutant albumin (R410A/Y411A/K525A), in which three of five high affinity fatty acid-binding sites were substituted for alanine and whose expression failed to sequester RA, did not inhibit HSC activation [20, 23].

Previous studies demonstrated that inflammatory mediators play a role in HSC activation [43], but we showed in this study a direct inhibitory interaction between the IL-1β and TGF-β signaling in HSCs. As albumin is expressed only in pre-activated HSCs and inhibits HSC activation, it may be used as a reliable marker for pre-activated HSCs.

Retinoid-storing stellate cells also exist in extrahepatic organs such as pancreas, kidney, spleen, intestine and lung [44]. These cells show striking similarities in morphology and perivascular location, which suggests that activated stellate cells may contribute to the myofibroblast cells seen in the fibrotic extrahepatic tissues [6]. As intravenously injected R-III was detected also in extrahepatic organs such as lung, pancreas, kidney and intestine [21], we are investigating whether R-III reduces extrahepatic fibrosis. In the present study, we elucidated the mode of action of albumin/R-III, and this study suggests that R-III, designed for stellate cell-targeting, may be a novel anti-fibrotic drug candidate.

## Materials and methods

### Materials

Male BALB/c mice (9 weeks of age) were purchased from Central Lab Animal, Inc. (Seoul, Korea) and maintained in proper condition. Animal protocols were approved by the Institutional Animal Care and Use Committee (IACUC) in Korea University. Mouse R-III fusion protein was prepared as previously described [23]. Citral, IL-1 receptor antagonist, (5Z)-7-Oxozeaenol, SB203580 and SP600125 were purchased from Sigma-Aldrich (St. Louis, MO, USA) and AGN193109 purchased from Santa Cruz Biotechnology (Dallas, TX, USA).

### Isolation of Mouse hepatic stellate cells (HSCs)

Mouse HSCs were isolated from wild-type male BALB/c mice (14 weeks old) as previous described [45] with some modifications. Briefly, liver was *in situ* perfused with phosphate-buffered saline (PBS) and then Gey’s balanced salt solution (GBSS) supplemented with collagenase, pronase (Sigma-Aldrich, St. Louis, MO, USA) and DNase (MP Biomedicals, Santa Ana, CA, USA). The perfused liver was dissected and the gall bladder and connective tissue attached to the liver were removed. The liver cell suspensions were further digested for 12 min in a 37°C water bath in GBSS supplemented with collagenase, pronase and DNase. The cells were then washed and centrifuged in a 13.4% Nycodenz gradient at 1400 g for 20 min without brake. The interface containing enriched HSC was taken, washed by GBSS, and HSCs were cultured in Dulbecco’s Modified Eagle Medium (DMEM) supplemented with 10% fetal bovine serum (FBS). The purity of stellate cells was assessed by microscopic observation and western blotting using anti-tyrosine aminotransferase antibody. The isolated cells were permitted to activate 4 days in culture and were then trypsinized and plated. Activation status of HSCs was assessed by increased expression of α-SMA and morphological changes.

### Quantitative Real-Time PCR

Total RNA was prepared using TRIzol (Ambion, Austin, TX, USA) and cDNA synthesized from total RNA. Quantitative real-time PCR was performed using an ABI QuantStudio™ 3 Real-Time PCR System. To control for variations in the reactions, PCR products were normalized against glyceraldehyde 3-phosphate dehydrogenase (GAPDH). The primers used were listed in Table S1.

### Western blot analysis

Cell lysates were prepared, and electrophoresis and immunoblotting were performed as described [23]. Primary antibodies used were α-SMA (Sigma-Aldrich #A2547, St. Louis, MO, USA), Smad2, p-Smad2 (S465,467), Smad3, p-Smad3 (S423,425), TAK1, p-TAK1 (S412), p38, p-p38, JNK, p-JNK, α-tubulin (#5339, 3101, 9523, 9520, 4505, 9339, 9212, 9211, 9252, 9255, 2125; Cell Signaling Technology, Beverly, MA, USA), p-Smad3 (T179) (Merck #ABS47, Darmstadt, Germany), p-Smad3 (S208), p-TAK1 (T184,187) (#PA5-38521, MA5-15073; Thermo Fisher Scientific, Waltham, MA, USA), PPAR-γ, RARα (#ab41928, ab41934; Abcam, Cambridge, UK), RXR (Santa Cruz Biotechnology #sc-774, Dallas, TX, USA).

### Immunoprecipitation

Cells were washed in cold PBS and placed in RIPA buffer containing 50 mM Tris pH8.0, 150 mM NaCl, 1% NP-40, 0.5% sodium deoxycholate with one tablet of Protease Inhibitor (Roche Life Sciences, Indianapolis, IN) per 50 ml of buffer. Equal amounts of protein were immunoprecipitated with anti-RXR polyclonal antibody (Santa Cruz Biotechnology #sc-774) for 4 hours at 4°C, followed by overnight incubation with 20 µl protein A/G PLUG-Agarose beads (Santa Cruz Biotechnology #sc-2003) at 4°C. The beads were then washed, boiled, and the supernatants were used to immunoblot with antibodies against PPAR-γ (Abcam #ab41928), RARα (Abcam #ab41934) and RXR (Santa Cruz Biotechnology #sc-774). All blots were performed in two independent experiments.

### Immunofluorescence

HSCs were plated onto glass coverslips coated with gelatin. Samples were fixed in 4% paraformaldehyde, permeabilized, blocked, and incubated with antibodies against p65 (Cell Signaling Technology #8242), Smad2 and Smad3 (Cell Signaling Technology #5339, #9523) overnight at 4°C in a moist chamber, followed by Alexa Fluor® 594–conjugated secondary antibody. Cells then were stained with DAPI, washed with PBS and mounted onto slides. Stained cells were visualised on a Leica DM6000 B fluorescence microscope (Leica Microsystem, Buffalo Grove, IL).

### RNA-seq

Detailed experimental procedure for RNA preparation and RNA-seq is provided in Supplementary information.

### Statistical analysis

Results are expressed as mean ± standard deviation (SD). Paired statistical analysis was done using t tests. Comparisons were considered significant at p,0.05 and p values were two tailed.

## Acknowledgements

We thank Ms. Mihee Yoon and M.S. U Suk Jung for the technical assistance, and Dr. Young-Sik Kim for encouragement and discussion. This work was supported by a grant from National Research Foundation of Korea (NRF) funded by the Ministry of Science and ICT (NRF-2017M3A9C8031617).

## Materials and Methods

### mRNA-Seq Data

In order to construct cDNA libraries with the TruSeq Stranded mRNA LT Sample Prep Kit, total RNA was extracted from HSCs. The protocol consisted of polyA-selected RNA extraction, RNA fragmentation, random hexamer primed reverse transcription and 100nt paired-end sequencing by Illumina NovaSeq 6000. The libraries were quantified using qPCR according to the qPCR Quantification Protocol Guide and qualified using an Agilent Technologies 2100 Bioanalyzer. We preprocessed the raw reads from the sequencer to remove low quality and adapter sequence before analysis and aligned the processed reads to the *Mus musculus (mm10)* using HISAT v2.1.0 (1). HISAT utilizes two types of indexes for alignment (a global, whole-genome index and tens of thousands of small local indexes). These two types’ indexes are constructed using the same BWT (Burrows–Wheeler transform)/a graph FM index (GFM) as Bowtie2. Because of its use of these efficient data structures and algorithms, HISAT generates spliced alignments several times faster than Bowtie and BWA widely used. The reference genome sequence of *Mus musculus (mm10)* and annotation data were downloaded from the NCBI. And then, transcript assembly of known transcripts, novel transcripts, and alternative splicing transcripts was processed by StringTie v1.3.4d (2, 3). Base on the result of that, expression abundance of transcript and gene were calculated as read count or FPKM value (Fragments Per Kilobase of exon per Million fragments mapped) per sample. The expression profiles are used to do additional analysis such as DEG (Differentially Expressed Genes). In groups with different conditions, differentially expressed genes or transcripts can be filtered through statistical hypothesis testing.

For variant calling of RNA-Seq data, trimmed reads were aligned to *Mus musculus (mm10)* with STAR (4) and then duplications were marked and discarded using Picard MarkDuplicate (http://broadinstitute.github.io/picard/). Afterwards, aligned reads that can be used in analysis are created through Split ‘N’ Trim, mapping quality reassignment, indel realignment, and base recalibration process. The aligned reads are used for variant calling with HaplotypeCaller module of GATK v2015.1-3.4.0-1-ga5ca3fc (5). Variant filtering for each sample was performed base on Fisher Strand values(FS >30.0) and Qual By Depth values(QC < 2.0) in the VariantFiltration module of GATK.

### Statistical analysis of gene expression level

The relative abundances of gene were measured in read count using StringTie. We performed the statistical analysis to find differentially expressed genes using the estimates of abundances for each gene in samples. Genes with one more than zeroed read count values in the samples were excluded. Filtered data were log2-transformed and subjected to TMM Normalization. Statistical significance of the differential expression data was determined using edgeR exactTest (6) and fold change in which the null hypothesis was that no difference exists among groups. False discovery rate (FDR) was controlled by adjusting p value using Benjamini-Hochberg algorithm. For DEG set, hierarchical clustering analysis was performed using complete linkage and Euclidean distance as a measure of similarity. Gene-enrichment analysis and KEGG pathway analysis for DEGs were also performed based on Gene Ontology (http://geneontology.org/) and KEGG pathway (https://www.genome.jp/kegg/) database respectively.

### Multidimensional scaling

We used multidimensional scaling (MDS) method to visualize the similarities among samples. MDS is one of the methods that convert the structure in similarity matrix to a simple geometrical picture as scatter plots. The larger the dissimilarity between 2 samples, the further apart the points representing the experiments in the picture should be. We applied to the Euclidean distance as the measure of the dissimilarity.

### Hierarchical clustering

Hierarchical clustering analysis also was performed using complete linkage and Euclidean distance as a measure of similarity to display the expression patterns of differentially expressed transcripts which are satisfied with |fold change|≥2 and raw p <0.05. All data analysis and visualization of differentially expressed genes was conducted using R 3.5.2 (www.r-project.org).

---

**FIG. S1.**
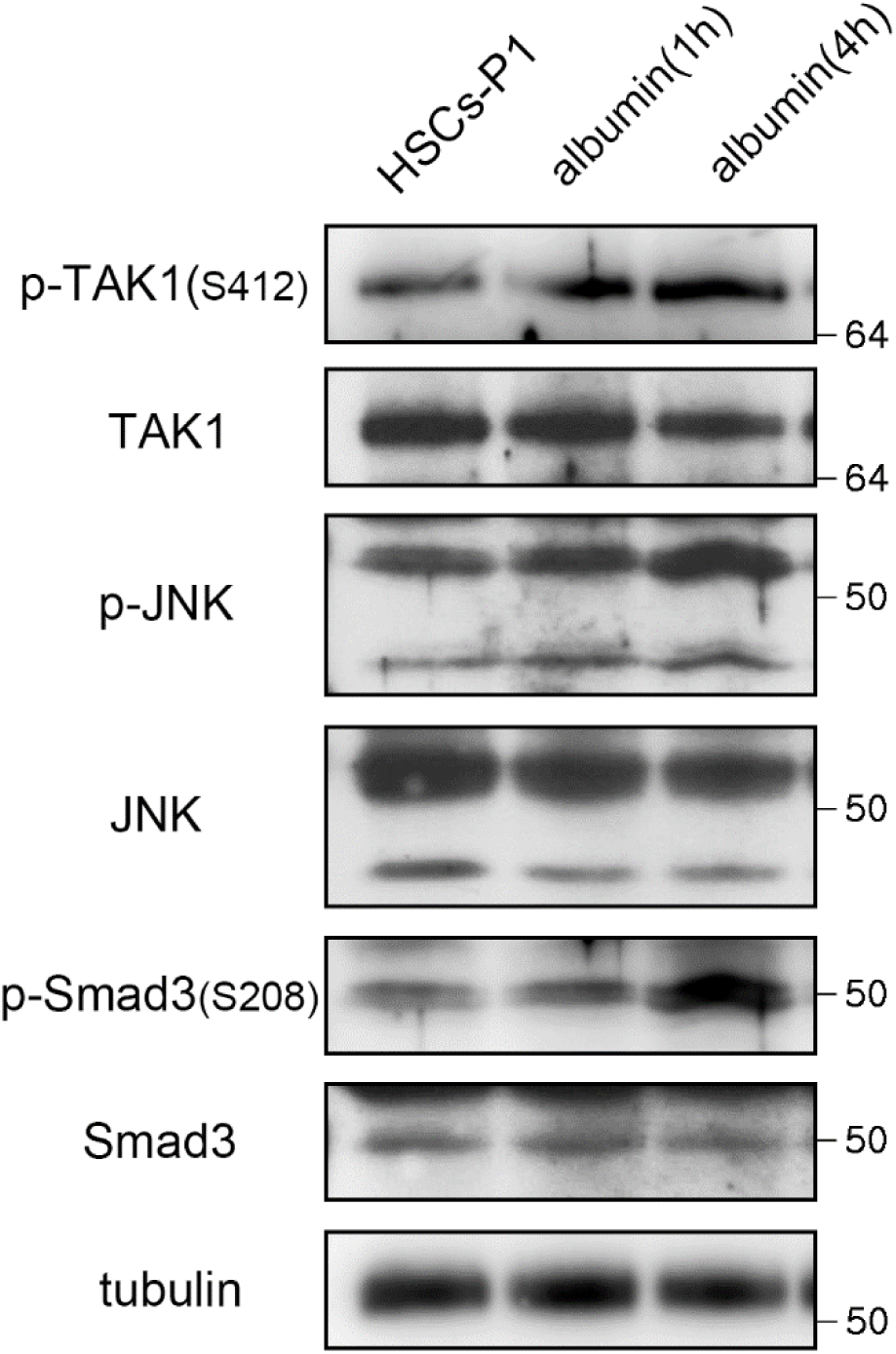
HSCs-P1 were transfected with albumin expression vector and cell lysates were analyzed by western blotting. The Western blots are representative of three independent experiments from separate cell preparations. α-tubulin was used as a loading control.

**Table S1.**
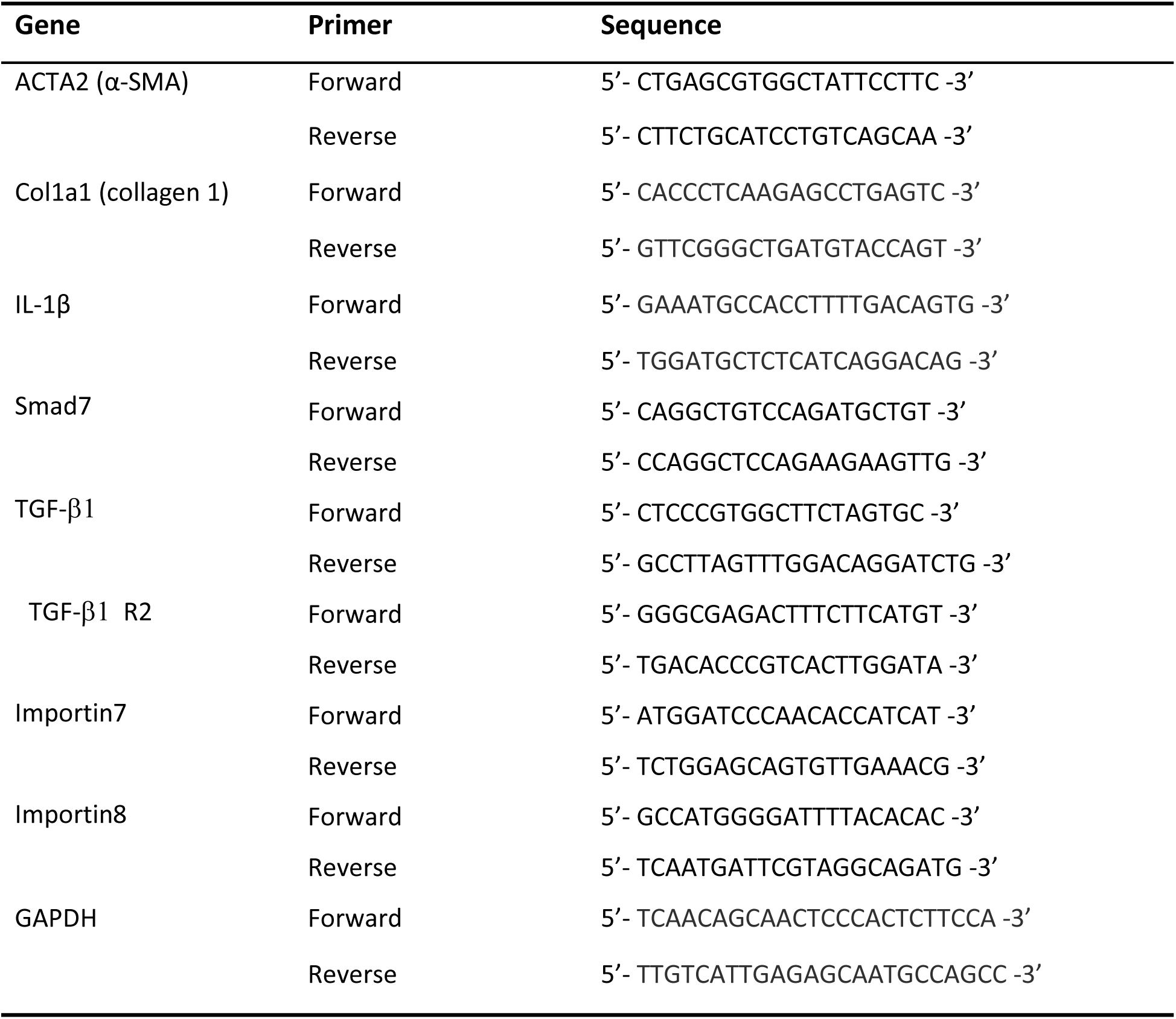
Primers used for real-time PCR.

